# Identification of histological features to predict MUC2 expression in colon cancer tissues

**DOI:** 10.1101/584292

**Authors:** Preethi Periyakoil, Michael F. Clarke, Debashis Sahoo

## Abstract

Colorectal cancer (CRC) is the third-most common form of cancer among Americans. Like normal colon tissue, CRC cells are sustained by a subpopulation of “stem cells” that possess the ability to self-renew and differentiate into more specialized cancer cell types. In normal colon tissue, the enterocytes, goblet cells and other epithelial cells in the mucosa region have distinct morphologies that distinguish them from the other cells in the lamina propria, muscularis mucosa, and submucosa. However, in a tumor, the morphology of the cancer cells varies dramatically. Cancer cells that express genes specific to goblet cells significantly differ in shape and size compared to their normal counterparts. Even though a large number of hematoxylin and eosin (H&E)-stained sections and the corresponding RNA sequencing (RNASeq) data from CRC are available from The Cancer Genome Atlas (TCGA), prediction of gene expression patterns from tissue histological features has not been attempted yet. In this manuscript, we identified histological features that are strongly associated with MUC2 expression patterns in a tumor. Specifically, we show that large nuclear area is associated with MUC2-high tumors (p < 0.001). This discovery provides insight into cancer biology and tumor histology and demonstrates that it may be possible to predict certain gene expressions from histological features.

## Introduction

Colorectal cancer (CRC) is the second leading cause of cancer-related deaths in the United States, in addition to being the third most common form of cancer.^1,2^ Approximately, 140,250 colon cancer cases expected to occur in the United States in 2019, with a projected death rate of 50,630 deaths.^1^ At the cellular level, CRC is typically diagnosed using hematoxylin and eosin (H&E), which highlight different tissue structures such as cell nuclei and cytoplasm in a tissue section.^3^ A pathologist manually examines the H&E stains to determine the cancer diagnosis, which is the current gold standard. Immunohistochemical analysis on the tissue section is performed to examine protein expression patterns at a single cell resolution.^4,5^ In a normal tissue sample, expression patterns of cell-type-specific markers are usually correlated with distinct morphology because differentiated cells, such as enterocytes and goblets cells, have distinct shapes. A typical histopathological image of CRC tissue contains four tissue components: lumen, cytoplasm, epithelial cells, and stroma (connective tissue, blood vessels, nervous tissue, etc.). However, it is hard to distinguish the different cell types within the epithelium in a cancer tissue sample. The epithelial cells in CRC come in diverse shapes and sizes. A recent review of mucin expression in CRC demonstrates this diversity comprehensively.^6^

Previous studies have observed that mucinous histology of CRC is strongly associated with overexpression of MUC2.^7^ Research on nuclear morphology and MUC2 expression has been limited. Cancer cells with signet-ring characteristics, such as clear cytoplasm and abundant intracytoplasmic mucins that compress the nuclei and cause them to be eccentrically located are characteristic of MUC2 overexpression. Knowledge of specific nuclear features may associated with MUC2 expression may help with devising ways to detect colon cancer in its early stages. Currently, we rely on simple nuclear segmentation techniques to identify nuclear morphology.^8^ By performing analysis on high-resolution images of H&E stained tissues and the corresponding RNA sequencing (RNASeq) data available from The Cancer Genome Atlas (TCGA)^9^, we sought to determine if there were specific nuclear features that are strongly associated with MUC2 expression.

## Methods

### Data Collection

Whole slide images (n = 897) and RNASeq data (n = 698) were downloaded from the TCGA website for colorectal cancer (COAD + READ). The RNASeq data was explored using a scatter plot between *KRT20* and *MUC2* TPM values (Fig. 1B). The expression values of *KRT20* and *MUC2* were ordinally arranged from low to high, and a rising step function was computed to define a threshold by the StepMiner algorithm in the TCGA dataset^10^. Expression levels above the assigned threshold for a gene were classified as “high”, and expression levels below this assigned threshold were classified as “low”. We identified 61 high-quality tissue images from 45 patients corresponding to 23 MUC2-low CRC, 28 MUC2-high CRC, 10 MUC2-high normal colon tissue. 61 TPM (transcripts per million) values corresponding to these 45 patients were obtained, including 23 MUC2-low CRC, 27 MUC2-high CRC, and 7 MUC2-high normal colon tissue. TPM values were not available for three normal colon tissues among the whole slide tissue images. However, since normal colon tissues are MUC2-high, we labeled these three tissues as MUC2-high samples. The Cancer Digital Slide Archive web interface was also used to explore the tissue images,^11^ and a proprietary web interface was developed (code available from authors upon request) to annotate the cancer regions in the tissue images.

**Fig. 1:**
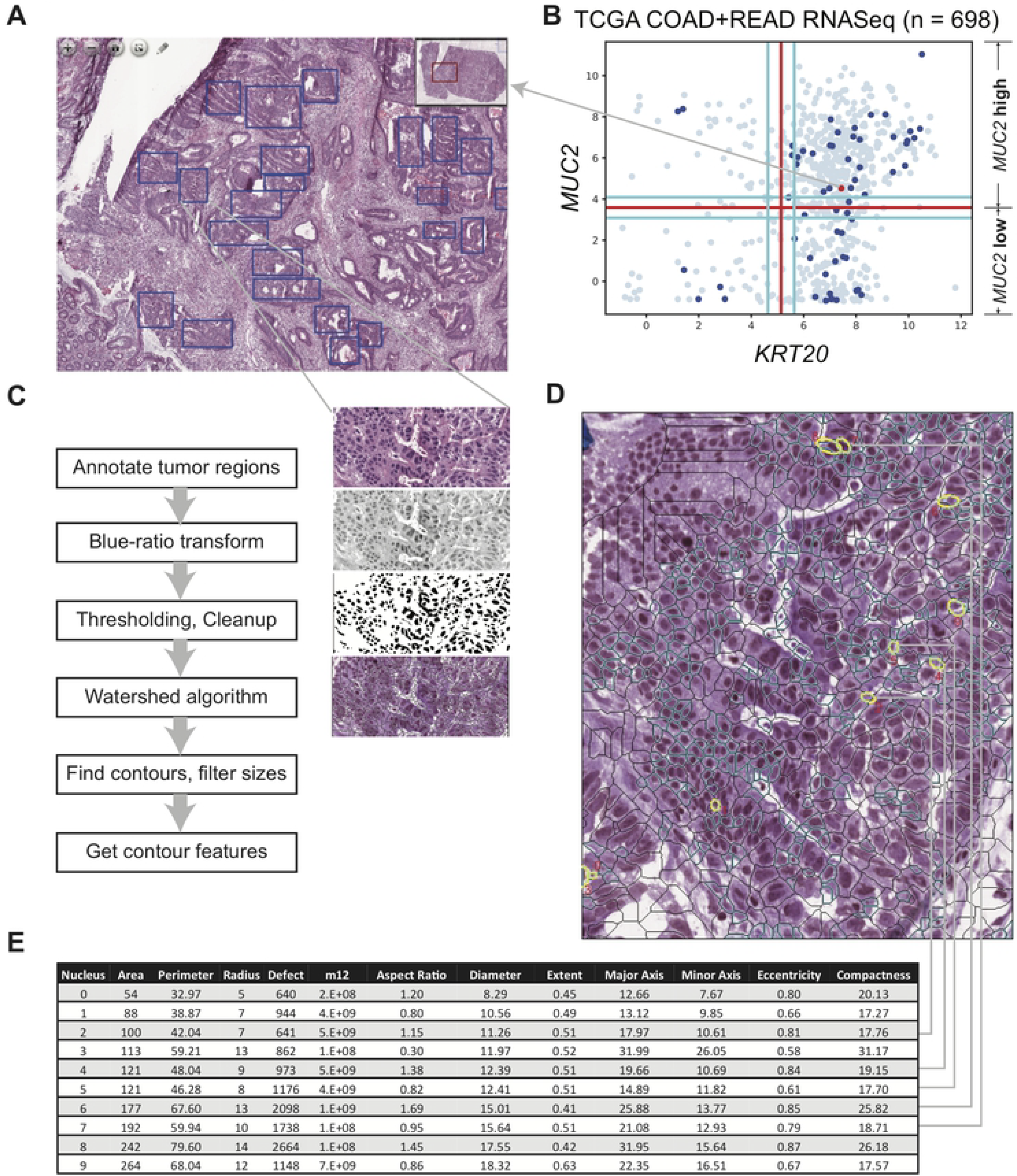
Tissue image analysis workflow. (A) Annotation and isolation of cancer regions (blue rectangles) in brightfield images of colorectal cancer (CRC) tissue cross-sections. (B) Scatter plot between *KRT20* and *MUC2* gene expression levels in the TCGA RNASeq dataset (n = 698). Red lines are StepMiner thresholds used to convert RAW data into Boolean high/low values. Cyan lines are noise margin around StepMiner threshold. Blue and red points (n = 61, 45 patients) refer to samples for which a whole slide tissue section image was annotated. Red point (n = 1) corresponds to the tissue shown in panel A. Gray points (n = 637) are other RNASeq experiments for which tissue image annotations were not available. (C) Individual steps of the image analysis workflow and illustrations (image transformations for each step) of each individual step of the nuclei segmentation process. (D) Results of the nuclear segmentation and ten randomly selected nuclei modeled as ellipses. (E) Table of the 13 selected computational features of the ten previously selected nuclei.

### Image Annotation

Each colon cancer tissue image contained several regions of cancer cell clusters, as well as of stromal cells, fibroblasts, muscle tissue, and necrotic tissue. The cancer regions, which were the specific focus of the study, were cropped and isolated and verified independently by two experts in cancer stem cell biology (Dr. Michael Clarke, Stanford University; and Dr. Piero Dalerba, Columbia University), who were blinded to each other. Each of these isolated regions, which will henceforth be referred to as annotations, was saved as a separate image file and labeled as MUC2-high (+1) or MUC2-low (−1), depending on the image from which it was cropped (which was based on the corresponding patient ID). From a total set of 897 images, 61 were randomly selected for the training set; these images were annotated and used for training and cross-validations. These images were manually annotated, and a total of 2336 annotations were created for image analysis.

### Image Analysis

For each annotation, OpenCV^12^ (an open-source computer vision software package) was used to segment the nuclei. Traditional computer-vision-based approaches follow four main steps of nuclei segmentation process: image enhancement, thresholding, feature extraction and refinements. The Blue Ratio transformation was used to enhance the nuclear staining while attenuating the background staining.^8^ The Blue Ratio transformation is performed using the following formula: 100* B / (1 + R + G) * 256 / (1 + R + G + B). Standard Otsu thresholding was performed to transform the grayscale transformed image into a binary image. To remove the artifacts of thresholding, the holes were filled and binary opening and closing morphological operations were performed.^13^ Connected components were discovered to form islands, and the watershed algorithm^14^ was performed to separate these islands. The shapes of the islands were detected using the standard OpenCV contour-finding algorithm, and these contours were used as the segmented nuclei.

### Filtering Nuclei Segments

An aggressive filtering strategy was utilized to minimize any computer vision-based nuclei segmentation process errors including those from fragmented nuclei and joined nuclei. At this point, it was apparent that some of the nuclei were joined to form large segments and other nuclei were fragmented. To remove the fused-nuclei segments, the nuclei were sorted based on their area from small to large, and the sum of the areas was computed. All nuclei with areas larger than half of the sum were removed because they were likely fused nuclei.

### Feature Analysis

Each nucleus was modeled as an ellipse because the nuclei in CRC tissues appeared elliptical in shape, and a set of numerical features was obtained for each individual nucleus. These features included: area, moments (of which there were 24), aspect ratio, perimeter, major axis, minor axis, radius, equivalent diameter, extent, defect, compactness, and eccentricity. This procedure led to the creation of approximately 1.5 million data points, one for each nucleus. These feature values were compared to the MUC2 expression patterns from the RNASeq data to discover potential relationships.

Very little correlation was observed between the nuclear features and the MUC2 TPM values. Thus, histograms were computed using the values for every individual feature between MUC2-high and MUC2-low tissue samples, and these histograms were compared with one another. The histograms showed that three specific features had predictive capabilities. For each of these three features, the fraction of the logarithmic values that were above a certain threshold specific to each feature were calculated for each image, and these fractions were statistically analyzed using the Welch Two Sample t-test. Finally, box plots were created to visualize the large value fractions for these features for MUC2-high and MUC2-low images across the entire data set.

### Statistical Analysis

All statistical analysis were performed in R, version 3.0.2 (2013-09-25) and in python 2.7, using scipy.stats.ttest_ind (SciPy v0.19.0). All tests were 2-sided at the P=0.05 level.

## Results

### Tissue histological features are compared with gene expression

Figure 1 depicts the process of annotating the images, segmenting the nuclei, and obtaining the numerical features for each nucleus. We collected tissue images from the TCGA website and manually annotated the cancer regions (Fig. 1A) and assembled TCGA RNASeq dataset to examine MUC2 gene expression patterns (Fig. 1B). Figure 1A shows a CRC tissue image that has high MUC2 gene expression levels, and the annotations of the cancer regions are depicted as blue rectangles. Figure 1C demonstrates the image transformations during individual steps of the nuclei segmentation. The final results of the nuclear segmentation, including the removal of the large fused nuclei, are shown in Figure 1D (cyan contours). Ten randomly selected nuclei were chosen to demonstrate the computational features (Fig. 1D, yellow ellipses). OpenCV was used to model the nuclei as ellipses and to obtain specific numerical features of each nucleus. Thirteen selected contour features out of 35 standard features are displayed in a table form to show the relationship between the feature values and the shape of the nuclei (Fig. 1E). These feature values were analyzed further to find relationships between the CRC tissue morphology and the expression levels of MUC2.

### Feature area, equivalent diameter, and m12 are associated with MUC2 expression in selected samples

To check the relationship between nuclei features and the MUC2 TPM values from the RNASeq experiments, we selected three MUC2-high samples, including a normal tissue sample and two MUC2-low CRC samples (Fig. 2A, 2B). A normal colon tissue sample is always MUC2-high because of the large number of goblet cells present, but it is not always clear from tissue images if a CRC sample is MUC2-high or low. A pooled nuclei data set was created from the three MUC2-high slide images (23993 nuclei) and two MUC2-low slide images (58195 nuclei), which led to a total of 82188 data points (Fig. 2C). No significant differences were observed between the average feature values between MUC2-high and MUC2-low groups.

**Fig. 2:**
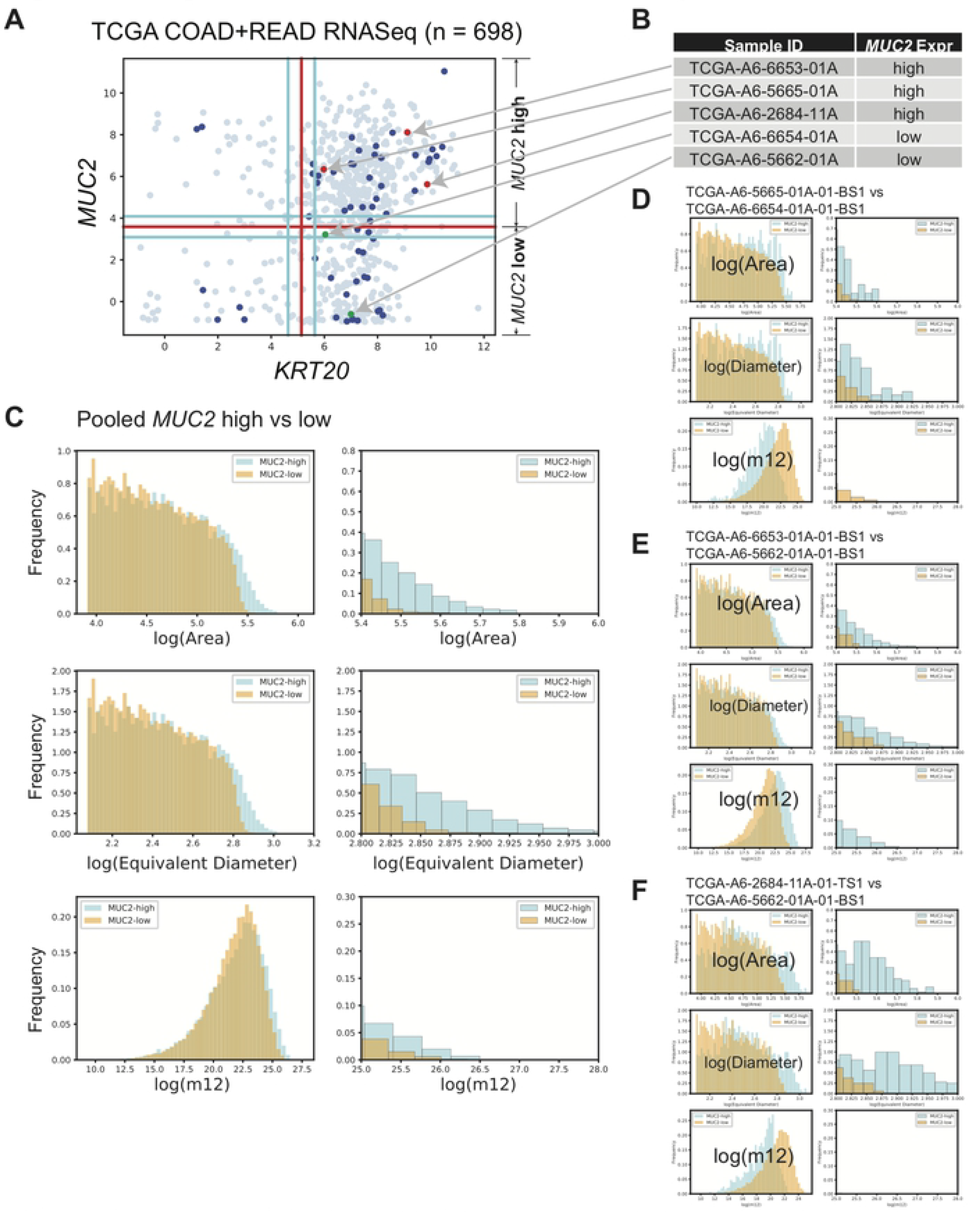
Image features associated with *MUC2* expression. (A) Scatter plot between *KRT20* and *MUC2* gene expression levels in TCGA RNASeq dataset (n = 698, 56 blue = tissue image annotation available, 3 red = selected *MUC2* high samples, 2 green = selected *MUC2* low samples, 637 gray= othersamples). (B) Three selected *MUC2* high (one normal colon tissue) and two selected *MUC2* low samples for image analyses and their corresponding points on the scatter plot shown in panel A using red and green color respectively. (C) Histograms of the area, equivalent diameter, and m12 image moment for the pooled segmented nuclei from the three selected *MUC2* high tissues compared to the two selected *MUC2* low tissues. Full histograms on the left, zoomed-in right tails on the right. (D, E) Comparison of histograms between one *MUC2* high with a *MUC2* low CRC sample. (F) Comparison of histograms between a normal colon tissue (positive control for *MUC2* high) with a *MUC2* low CRC sample.

With this pooled data set, a histogram was plotted for each feature generated by OpenCV to reveal any differences between MUC2-high and MUC2-low nuclei. No significant differences between MUC2-high and MUC2-low were observed in the distributions for any individual feature in a linear scale, so they were again plotted on a logarithmic scale. The following three features were found to have differences in the distributions between MUC2-high and MUC2-low (Fig. 2C): area, equivalent diameter, and m12 (m12 is one of the 24 image moments that were collected as individual features). The right tails of the histograms show that MUC2-high tissue has more nuclei that have higher values for all three features. Because the pooled data set had more MUC2-low nuclei than MUC2-high nuclei, it was surprising to find that more MUC2-high nuclei had higher values for these four features. Figure 2C right side shows a closer view of the right tails of each of the histograms shown in the left.

The results shown in the histograms led to the hypothesis that MUC2-high tissue may have more nuclei with larger area, equivalent diameter, and m12 than MUC2-low tissue do. To test if this hypothesis holds for individual images instead of the sample dataset, one MUC2-high sample was compared with one MUC2-low sample (Fig. 2D-F). The comparison between TCGA-A6-5665-01A-01-BS1 (MUC2-high) and TCGA-A6-6654-01A-01-BS1 (MUC2-low) revealed that only log(Area) and log(Equivalent Diameter) have a longer right tail in the MUC2-high case (Fig. 2D). However, the comparison between TCGA-A6-6653-01A-01-BS1 (MUC2-high) and TCGA-A6-5662-01A-01-BS1 (MUC2-low) reported similar results to those of the pooled dataset: area, equivalent diameter, and m12 shows longer right tail in MUC2-high sample (Fig. 2E). The normal tissue sample TCGA-A6-2684-11A-01-TS1 was compared with a MUC2-low CRC sample TCGA-A6-5662-01A-01-BS1, and the long right tail was only observed for area and equivalent diameter (Fig. 2F).

### Feature area and equivalent diameter are significantly associated with MUC2 expression in TCGA dataset

To test the hypothesis that area, equivalent diameter, and m12 may be associated with MUC2 expression patterns, analysis was performed on additional tissue samples from the TCGA dataset. 23 MUC2-low samples and 38 MUC2-high samples (28 CRC + 10 normal) from 45 patients with high quality tissue images were assembled, along with three normal tissue samples were not profiled in the RNASeq data (these samples were considered MUC2-high). 61 RNASeq experiments (54 CRC + 7 normal) correspond to these 45 patients (Fig 3A, 23 blue MUC2-low, 34 red MUC2-high, 2 cyan MUC2-low, 2 green, MUC2-low). Two patients were found whose tumor samples (Fig 3A, 2 cyan, 2 green) were on both side of the MUC2 threshold; these samples were treated as MUC2-low as they were boundary cases. The 61 tissue images (23 MUC2-low + 38 MUC2-high) were analyzed and the nuclei identified and segmented.

**Fig. 3:**
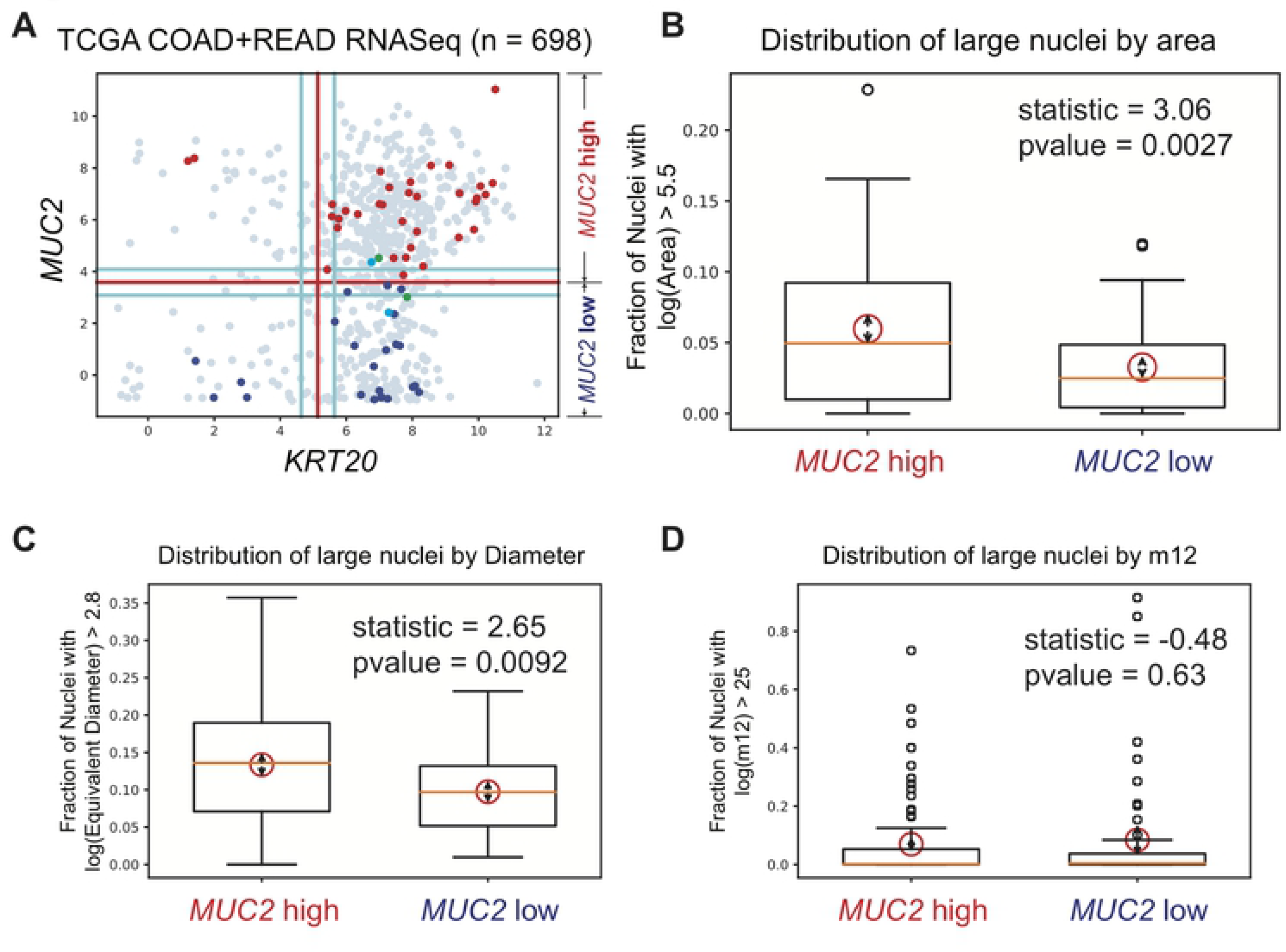
Validation of *MUC2* associated features. (A) Scatter plot between *KRT20* and *MUC2* gene expression levels in TCGA RNASeq dataset (n = 698, 23 blue = annotated MUC2 low, 34 red = annotated *MUC2* high samples, 2 green and 2 cyan = two CRC samples from the same patients in the opposite side of the threshold, 637 gray= other samples). (B, C, D) Boxplots of *MUC2* high and *MUC2* low samples. Center of the red circle represent the mean values and the arrow lengths represent the 95% confidence interval. Comparison of the mean values are performed using a T-test (statistic and pvalue are shown). (B) Boxplots of the fraction of nuclei whose log(Area) > 5.5 between *MUC2* high and *MUC2* low samples (p = 0.0027). (C) Boxplots of the fraction of nuclei whose log(Equivalent Diameter) > 2.8 between *MUC2* high and *MUC2* low samples (p = 0.0092). (D) Boxplots of the fraction of nuclei whose log(m12) > 25 between *MUC2* high and *MUC2* low samples (p = 0.63).

For area, equivalent diameter, and m12, the fraction of the logarithmic values that were above a certain threshold were calculated for each image, and these fractions were statistically analyzed using the Welch Two Sample t-test. The results of the t-test are illustrated in Figure 3 as red circles with the center as the mean, the arrow length as 95% confidence interval on both sides, the test statistic value, and the p-value. The p-values shown in Figure 3B and 3C show that the results found for area and equivalent diameter were statistically significant, while those for m12 were not (Fig. 3D).

Box plots were created to visualize the large value fractions for area, equivalent diameter, and m12 across all MUC2-high and MUC2-low images in the data set (Figure 3B-D). The box plots show that the median values of area and equivalent diameter are higher for MUC2-high cells than for MUC2-low cells. Because MUC2 is usually identified through tissue staining, the morphology of nuclei that are positive for MUC2 is not well known. These results suggest that tissue containing high volumes of large nuclei (or nuclei with large equivalent diameters) may be associated with high levels of MUC2. Moments are a quantitative measure of the spatial distribution of a set of points, and for these images, these moments are based on pixel intensity. It is unknown how this information correlates with existing information about colon cancer nuclei, and thus further experimentation should be conducted to analyze this relationship.

## Discussion

The Cancer Genome Atlas (TCGA) has published thousands of whole slide images from CRC tissues, in addition to RNASeq and genomic data on the same tissues. Literature and the field of digital pathology have focused on analysis of the tissue images using novel computational tools.^15^ However, the relationship between image features and gene expression has not been pursued aggressively. In this study, we demonstrate a strong association between tissue image features and MUC2 expression patterns. We chose MUC2 as our gene of interest because it is expressed in goblet cells, which have a distinct morphology. It is easy to identify goblet cells in normal tissue; but it is difficult to do so in CRC tissue.

The Blue Ratio transform was found to be effective at segmenting nuclei, and recording the features of individual nuclei was expected to prevent loss of data. Histograms were created for each nuclear feature generated by OpenCV, for a small subset of the slide images that were obtained from TCGA. While most of the features were indistinguishable in MUC2-high and MUC2-low tissue, three features in particular (area, equivalent diameter, and m12) were found to have a higher percentage of large values in MUC2-high tissue than in MUC2-low tissue. Discovery of such significant morphological difference between MUC2-high and MUC2-low nuclei is extremely relevant to the field of digital pathology. Connecting morphological features with gene expression will provide insight into the disease phenotype and their correlation with outcome. Although these results are surprising, further research must be conducted to investigate the correlation between MUC2 expression and nuclear size. The relative levels of gene expression in a tissue cross-section are often identified by staining the tissue with an antibody specific to that gene, but such information is very difficult for humans to identify by eye in unstained brightfield images. The fact that a computer is able to relate a morphological feature in the tissue sample with relative gene expression is important step towards understanding CRC disease diversity.

Our study is limited by the consideration that only 61 slide images were analyzed, even though large volumes of data were produced from said images. Because the cells in cancer tissue are extremely heterogeneous in shape, size, and other morphological features, it is necessary to have a high volume of data that is all encompassing of these morphological features. Thus, more images must be annotated and added to the data set, and any computed results will thus be more accurate. The other limitation of this study is that cells expressing MUC2 are difficult to identify by eye, and thus the segmentation algorithms have not been trained on cells that are specifically known to express MUC2. Thus, this study must be conducted on a training set that consists of tissues specifically stained for MUC2.

Insight gained through this analysis is not only relevant to the field of cancer biology and tissue histology but also relevant to the field of bioinformatics, which provides tools for predicting gene expression directly from tissue H&E slides. Advanced nuclear segmentation techniques can be used in future to improve this analysis: CellProfiler,^16,17^ QuPath,^18^ Fiji^19^ and MANA.^20^ Deep learning approaches are ideally suited for this type of problems because of availability of large number of training and validation data. Doing so will enable the models to better identify features that are indicative of MUC2-high cells, and will thus make them more effective at predicting a tissue sample as MUC2-high or MUC2-low.

Image segmentation is an active area of research,^21^ and nuclear segmentation techniques have improved significantly.^15^ A search for “digital pathology” and “image analysis” on Pubmed revealed over 1500 articles just in the last decade,^15^ and an extensive collection of nuclear segmentation papers for histologic analysis have been published over the past 20 years.^22^ Several machine learning/deep learning-based approaches have been proposed to perform better than computer vision-based approaches.^21,23^ Histopathology images are ideally suited for interrogation via deep learning approaches as they attempt to use deep architectures to learn complex features from data in an unsupervised manner.^15^ Much progress has been made in the segmentation of glandular structure in a tissue image.^21,24–29^ A nuclear segmentation algorithm consisting of color deconvolution, k-means clustering, local adaptive thresholding, and cell separation was explored to extract cell nuclei features in a recent study,^3^ and previous studies have used active contour models to segment nuclei.^30–32^ It is important to replicate our results in the context of other nuclei segmentation approaches to study the robustness of the association.

## Data Access

TCGA

## Acknowledgements

This study was supported by following three institutions: The California Institute of Technology; Stanford University; University of California, San Diego. This work was supported by the National Institutes of Health (NIH) grant #R00-CA151673 to DS, 2017 Padres Pedal the Cause / Rady Children’s Hospital Translational PEDIATRIC Cancer Research Award to DS, 2017 Padres Pedal the Cause /C3 Collaborative Translational Cancer Research Award to DS.

